# Genomic signatures of bottleneck and founder effects in dingoes

**DOI:** 10.1101/2023.02.05.527211

**Authors:** Manoharan Kumar, Gabriel Conroy, Steven Ogbourne, Kylie Cairns, Liesbeth Borburgh, Sankar Subramanian

## Abstract

Dingoes arrived in Australia during the mid-Holocene and are the native top-order terrestrial predator on the mainland and some offshore islands. Although dingoes subsequently spread across the continent, the initial founding population(s) could have been small. To investigate this hypothesis, we examined the potential signatures of bottlenecks and founder effects in dingoes by sequencing the whole genomes of three dingoes and also obtaining the genome data from nine additional dingoes and 56 canines, including wolves, village dogs and breed dogs, and examined the signatures of bottlenecks and founder effects. We found that the nucleotide diversity of dingoes was low, and 36% less than highly inbred breed dogs and 3.3 times lower than wolves. The number of runs of homozygosity (RoH) segments in dingoes was 1.6 to 4.7 times higher than in other canines. Whilst examining deleterious mutational load, we observed that dingoes carried elevated ratios of nonsynonymous-to-synonymous diversities, significantly higher numbers of homozygous deleterious Single Nucleotide Variants (SNVs), and increased numbers of loss of function SNVs, compared to breed dogs, village dogs, and wolves. These results suggest dingoes experienced a severe bottleneck, potentially caused by the limited number of founding individuals. While many studies observe less diversity and a higher number of deleterious mutations in domesticated populations compared to their wild relatives, we observed the opposite – .i.e. wild dingoes have lower diversity and a greater number of harmful mutations than domesticated dogs. Our findings can be explained by bottlenecks and founder effects during the establishment of dingoes on mainland Australia. These findings highlight the need for conservation-based management of dingoes and need for wildlife managers to be cognisant of these findings when considering the use of lethal control measures across the landscape.

## Introduction

Dingoes arrived in Australia between 4,000 and 11,000 years ago, and their ancestors probably originated in Asia (Savolainen, et al. 2004; Cairns and Wilton 2016; Balme, et al. 2018; Bergstrom, et al. 2020; Zhang, et al. 2020). While their taxonomic status and evolutionary history remain contested, studies demonstrate their important ecological role as Australia’s largest terrestrial top-order predator (Johnson, et al. 2007; Wallach, et al. 2010; Letnic, et al. 2012) and also their cultural significance, particularly for First Nations Peoples (Corbett 2001; Archer-Lean, et al. 2015). Currently, dingo populations are found across the entire continent (excluding Tasmania), occupying vastly different habitats ranging from the mountainous Australian Alpine regions, tropical, subtropical, and temperate regions, the arid zones of Australia and offshore coastal barrier islands such as K’gari (Fraser Island). Dingo populations can vary widely in phenotypic features such as body weight, coat colour, and social behaviour (Corbett 2001; Koungoulos 2020; Tatler, et al. 2020; Cairns, et al. 2021). Despite their continent-wide presence and array of phenotypic variation, the size of the founding population of dingoes is likely to be small, given Australia’s geographic isolation, the narrow window for their arrival, and the potential that they were transported by humans. With the advent of cost-effective sequencing technologies, it is now possible to obtain the whole genome sequences and examine the genetic signatures of founder effects in the dingoes.

The nucleotide diversity of a population is determined by mutation rate (*μ*) and effective population size (*N*_*e*_) (Hartl 2007). Since the *μ* of a species is fairly constant over time, the reduction in nucleotide diversity is dictated by *N*_*e*_ (Kimura 1983). Therefore, measuring nucleotide diversity using whole genome data is a simple measure to detect the founder effects. The use of whole genome data also enables the measurement of lengths of long tracts of homozygous genotypes or runs of homozygosity (RoH) segments (Ceballos, et al. 2018). A number of previous studies showed that bottleneck and founder effects increase the number of long RoHs (Bortoluzzi, et al. 2020; Khan, et al. 2021; Sánchez-Barreiro F 2021; Tournebize, et al. 2022). Hence, estimating the lengths of RoH in dingoes and comparing them with those observed for other canines will potentially reveal whether the size of the founding dingo population was small. The third hallmark of the founder effect is the accumulation of deleterious genetic variations (Livingstone 1969; Evans 2015). The ratio of diversities at nonsynonymous and synonymous sites is routinely used to measure the deleterious mutational load in populations (Lohmueller, et al. 2008; Marsden, et al. 2016; Subramanian 2016; Makino, et al. 2018). These studies observed a much higher ratio in small populations than those in large populations, and the high ratio suggests a higher proportion of nonsynonymous Single Nucleotide Variants (SNVs). Reduction in population size was also known to increase the frequencies of rare deleterious variants, and hence the number of homozygous SNVs is expected to be high in bottlenecked populations. Indeed, significantly higher numbers of homozygous nonsynonymous SNVs are frequently reported in small or bottlenecked populations (Marsden, et al. 2016; Xie, et al. 2018; Subramanian 2021). Studies of bottlenecked populations also observe an abundance of homozygous SNVs that were predicted to be deleterious based on their functional consequences or evolutionary constraints. Therefore, the founder effect on dingo populations could be investigated by using whole genome data and measures that quantify the magnitude of the accumulation of deleterious variants in their genomes. Such analyses could provide insight into management concerns for this ecologically and culturally significant species that currently suffer from persecution in most Australian jurisdictions and are subject to widespread lethal control measures.

In our study, we investigated the genomic evidence for founder effects and bottlenecks in dingo populations by sequencing the whole genomes of three dingoes and obtaining the genome data of nine additional dingoes from mainland Australia. We also obtained genome data of breed dogs, village dogs, and wolves to give context to our analysis. We performed comparative analyses of these canine genomes and estimated various measures to capture the genetic diversity, runs of homozygosity, and accumulation of deleterious genetic variants.

## Materials and Methods

### Samples and whole-genome sequencing

The saliva samples from three dingoes were collected: one from Kimberley in Western Australia, one from western New South Wales, and a captive individual from the Alpine region of Southeast Australia. The samples were shipped to Novogene Co., Ltd. Beijing, China, for DNA extraction and sequencing. DNA extraction was performed by following standard operating protocols using saliva/tissue extraction kits. The library building and sequencing were performed following Illumina standard protocols. The genomic DNA was randomly sonicated into short fragments. Later, these fragments were end-repaired, added with A-Tail in the end, and ligated with Illumina adapter. Then the fragments only with adapters were PCR amplified, size selected, and purified for short read sequencing. Furthermore, the purified library was checked with Qubit and real-time PCR for quantification and a bioanalyzer for size distribution detection. Quantified libraries were sequenced on Illumina platforms, according to effective library concentration, to obtain 30 times coverage with 150 base pair paired-end reads.

### Whole-genome data collection

In addition to the three samples from this study, we obtained the raw sequence reads in *fastq* format for nine mainland dingoes from a previous study (Zhang, et al. 2020). The 12 samples come from the North (1), Northwest (2), Northeast (3), West (1), Central (1), and Southeast (4) regions of the Australian continent (See Supplementary Figure 1). We excluded one dingo genome from K’gari (Fraser Island) reported by Zhang et al (2020). A previous study showed a significant reduction in the heterozygosity of the K’gari-Fraser Island dingo population compared to the mainland dingoes (Conroy, et al. 2021). Therefore, we excluded the Island dingo in order to avoid the confounding effect of an additional bottleneck that could have occurred when the subset(s) of mainland dingoes became Island bound. In addition to dingoes, we obtained whole genome *fastq* data from 11 wolves, 13 village dogs, and 32 breed dogs (see Supplementary Table S1) from another study (Plassais, et al. 2019). The 32 dog breeds were chosen based on two criteria. First, we obtained dog breeds representing various geographical locations, including the Arctic, African, modern European, Asian and Middle eastern regions. The geographic locations of the breeds also reflect their distinct clades in the canine phylogeny (Vonholdt, et al. 2010). Second, out of 722 breed dogs from the previous study (Plassais, et al. 2019) we selected ten breeds having very high inbreeding coefficients. Since a very small number of founding individuals might have been used during breed formation, these breeds are the best candidates for comparison with dingoes. We also included the coyote genome obtained from the same study (Plassais, et al. 2019) to use it as the outgroup. The *fastq* sequences of these genomes were downloaded from the NCBI SRA database (https://www.ncbi.nlm.nih.gov/sra). Relevant SRA ID details can be found in (Table S1) supplementary information. These fastq reads were processed along with the newly sequenced three dingo samples as described below.

### Bioinformatics analyses

Illumina raw reads of three dingo samples were trimmed based on quality and adapter sequences using Trimmomatics-0.36 (Bolger, Lohse, & Usadel, 2014) (options: ILLUMINACLIP: adapter.fa:2:30:10 SLIDINGWINDOW:4:20 MINLEN:50). Similarly, we processed fastq reads of 68 samples of dingoes (11) and other canines (56) plus coyote obtained from published studies (Plassais, et al. 2019; Zhang, et al. 2020). Then, each sample’s clean reads were aligned to the canfam3 (https://hgdownload.soe.ucsc.edu/goldenPath/canFam3/bigZips/canFam3.fa.gz) reference genome using the bwa tool (Li & Durbin, 2009) (options: bwa mem -M). Mapped reads in the sequencing alignment mapped (SAM) format was converted to binary alignment/mapped (BAM) format using samtools version 1.31 (options: samtools view -@ 8 -b -F 4). Then, we sorted the aligned reads based on chromosomes in BAM format. The sorted reads were used for marking PCR duplicates using the Picard tool. Later, we generated genotypes for all sites using samtools (Li, 2011) (options: samtools mpileup -C 50 -uf | bcftools call -c -V indels -O z). Finally, we merged all position genotypes of 68 genomes using bcftools merge (par: -m all). After merging, we filtered biallelic variant sites using an in-house awk script. The final dataset had a total of 29,153,123 biallelic positions.

We carried out biallelic variant sites annotation using the *SNPeffect* tool (De Baets, et al. 2012) (options: java -jar snpEff.jar eff CanFam3) to obtain coding and non-coding sites. In addition, we also estimated the average read depth for each sample using the “depth” module of the software *samtool*. The total number of sites covered was calculated for each sample using an in-house script. The number of runs of homozygosity segments was estimated using the software *plink* (Purcell, et al. 2007b) with following parameter (--geno 0.01 --homozyg -- homozyg-window-het 0 --maf 0.05). To identify the ancestral state of the variants, we used the whole genome data of coyote to determine the direction of mutational change and to identify the derived alleles.

### Population genetics and statistical analyses

The software *plink* (Purcell, et al. 2007a) was used to perform Principal Component Analysis (PCA). The Maximum Likelihood-based method *Admixture* (Alexander, et al. 2009) was used to determine the admixture patterns of canine genomes. While we observed the lowest cross-validation error for K=2 (two ancestries), the canine populations were not clearly separated at this level. Therefore, we used K=4 to show the admixture patterns among canines as it separated the ancestries of dingoes, village dogs, breed dogs, and wolves. Higher K values (5 and 6) did not split the dingo populations further (but divided the breed dogs), suggesting more homogeneity among the dingoes used in this study. The inbreeding coefficients were calculated for the breed dogs using *plink*. This revealed a large variation in these estimates among breeds (0.12 - 0.92). Therefore, any mean estimate computed for breed dogs as one group will have confounding effects. To remove this bias, we grouped breed dogs into three categories based on their inbreeding coefficients. The High, Moderate, and Low inbred categories consist of breeds having inbreeding coefficient estimates of >0.4 (12), 0.4-0.2 (13), <0.2 (6), respectively. To identify deleterious missense alleles, we used the *SIFT* score (Ng and Henikoff 2003), and biallelic variants with a score <=0.05 were designated as ‘deleterious’. Loss of function (LoF) SNVs were identified based on the annotations “stop_lost”, “stop_gained”, “start_lost”, “splice_donor” and “splice_acceptor”. The numbers of synonymous and nonsynonymous SNVs were divided by their respective number of synonymous and nonsynonymous sites to obtain the diversities at these sites. The ratio of the two was used to determine the strength of purifying selection or the accumulation of deleterious nonsynonymous SNVs in each genome. The average number of LoF and deleterious SNVs per genome, along with the standard errors, were estimated for each genome. The significance between the mean counts was determined using the Z test, and the statistical significance was determined using the software Z to P (http://vassarstats.net/tabs_z.html). A Pearson correlation coefficient was used to determine the strength of the correlation. Furthermore, using the non-parametrical Spearman’s correlation also produced similar strength of correlation. The statistical significance of the correlation was determined by converting the correlation coefficient *r* to the normal deviation Z, and this was accomplished using the online software *r* to *P* (http://vassarstats.net/tabs_r.html).

## Results

### Nucleotide diversity based on the whole genomes of dingoes and other canines

In this study, we sequenced three dingo genomes and assembled the whole genome dataset from previously published dingoes, dog breeds, village dogs, and wolves (see Supplementary Table S1). Using this data, we performed principal component analysis to determine the level of population structure among the canines. This revealed three distinct clusters of dingoes, dogs, and wolves (Figure 1A. To investigate the genetic admixture, particularly for dingoes, we conducted an admixture analysis (Figure 1B). This showed one individual from Southeastern Australia had a small amount of admixture (14%) from dogs, which suggests that this may not be a ‘pure’ or unadmixed dingo. Therefore, we excluded this dingo from further analysis and estimated the genomic heterozygosity using the whole nuclear genomes of 11 dingoes and other canines. We found dingoes to have the lowest genomic nucleotide diversities compared to all other canines analysed, except for Norwegian Lunde Hund and Bull terrier dog breeds. We then grouped breed dogs into three categories based on their inbreeding coefficients (see methods). The mean diversity estimate of dingoes was significantly lower (*P* < 0.0001) than that of other canine groups (Figure 2 - inset). For instance, the genomic diversity of dingoes was 36% less than that of the highly inbred breed dogs. The mean estimates of the three dog breed groups (high, moderate, and low) vary significantly, and the diversity of village dogs was less than that of the low-inbred group. The mean diversity of wolves was found to be the highest of all canine groups, and it was 3.3 times higher than observed for dingoes.

**Figure 1.**
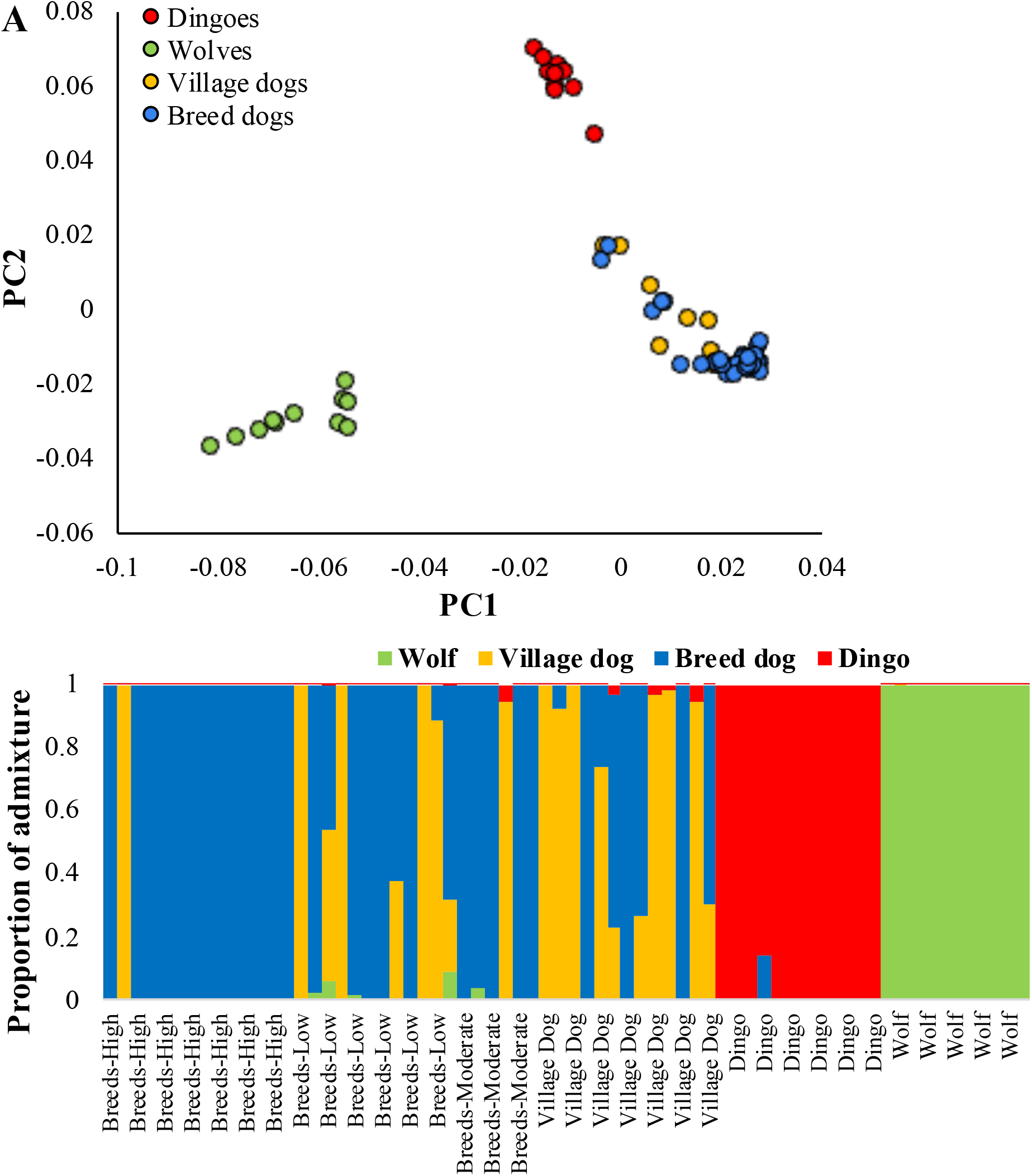
**(A)** The scatter graph showing the results from the principal component analysis (PCA) using the whole genomes of dogs, dingoes, and wolves. **(B)** The proportion of genetic admixture in each canine genome is shown. This analysis was performed assuming four ancestries for canines (K=4).

**Figure 2.**
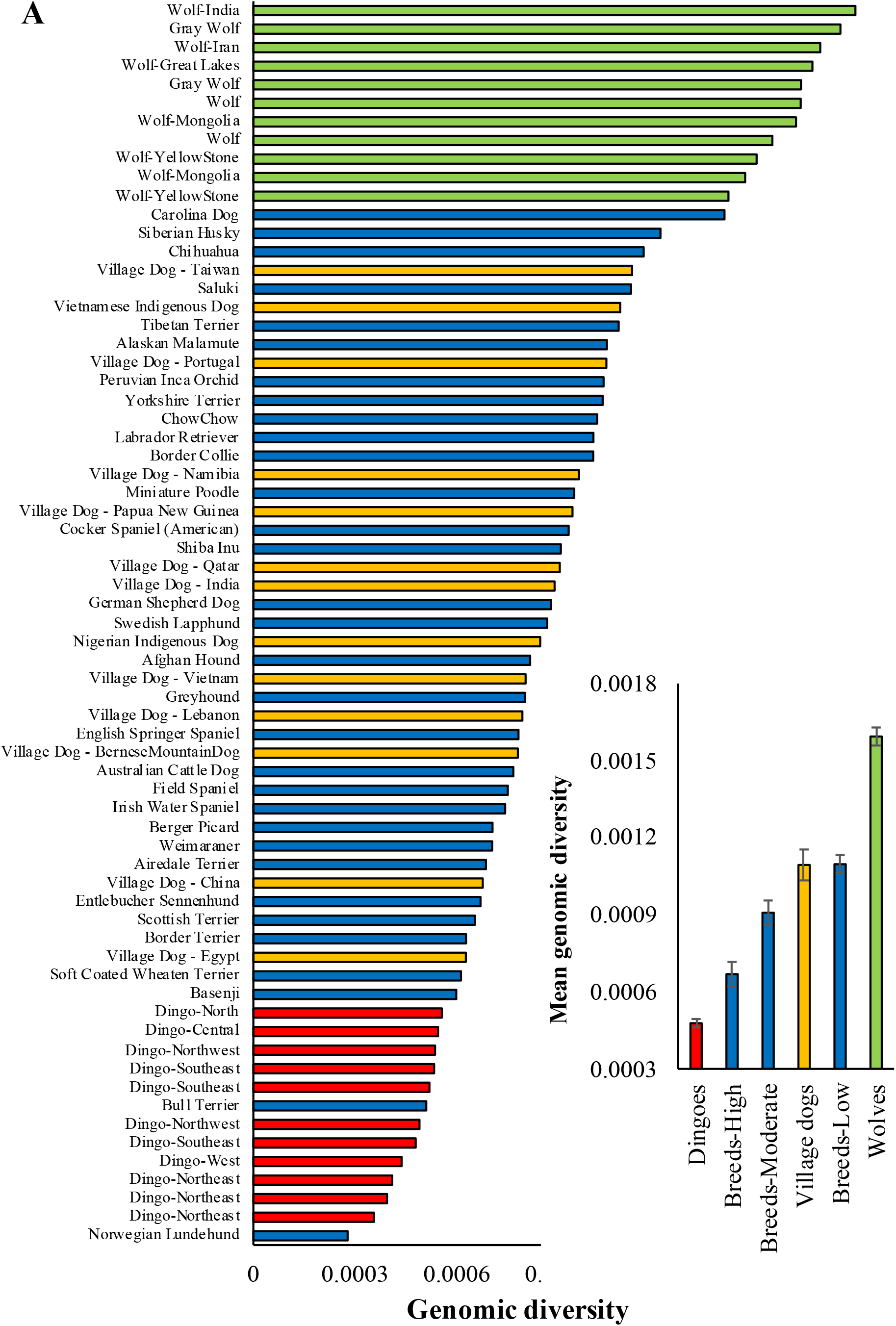
Genomic diversity estimates for dingoes and other canines. Nucleotide diversity was estimated using the whole genome sequences. The dog breeds were grouped into three categories based on their inbreeding coefficients – Highly inbred (>0.4), Moderately inbred (0.4-0.2), and lowly inbred (<0.2). **Inset**: Average genomic diversities computed for various canine groups. Error bars show the standard error of the mean.

### Runs of Homozygosity

Using the whole genome data, we next estimated the number of runs of homozygosity (RoH) segments. For this analysis, we used a threshold of 0.2Mb to define an RoH segment, and we also used a stringent criterion of not allowing any heterozygous SNVs within an RoH. This analysis revealed that dingoes have a much higher number of RoH segments than any other canine group (Figure 3). The mean number of RoH segments in dingoes was 2125, which was 62% higher than that estimated for the highly inbred breed dogs (1313), and the difference between the estimates was highly significant (*P* < 0.0001). The number of RoH segments per genome estimated for breed groups varied 1.6 times as the high and low inbred groups had 1313 and 814 segments, respectively. The average number of RoH segments in village dogs (605) and wolves (451) were four times smaller than that of dingoes.

**Figure 3.**
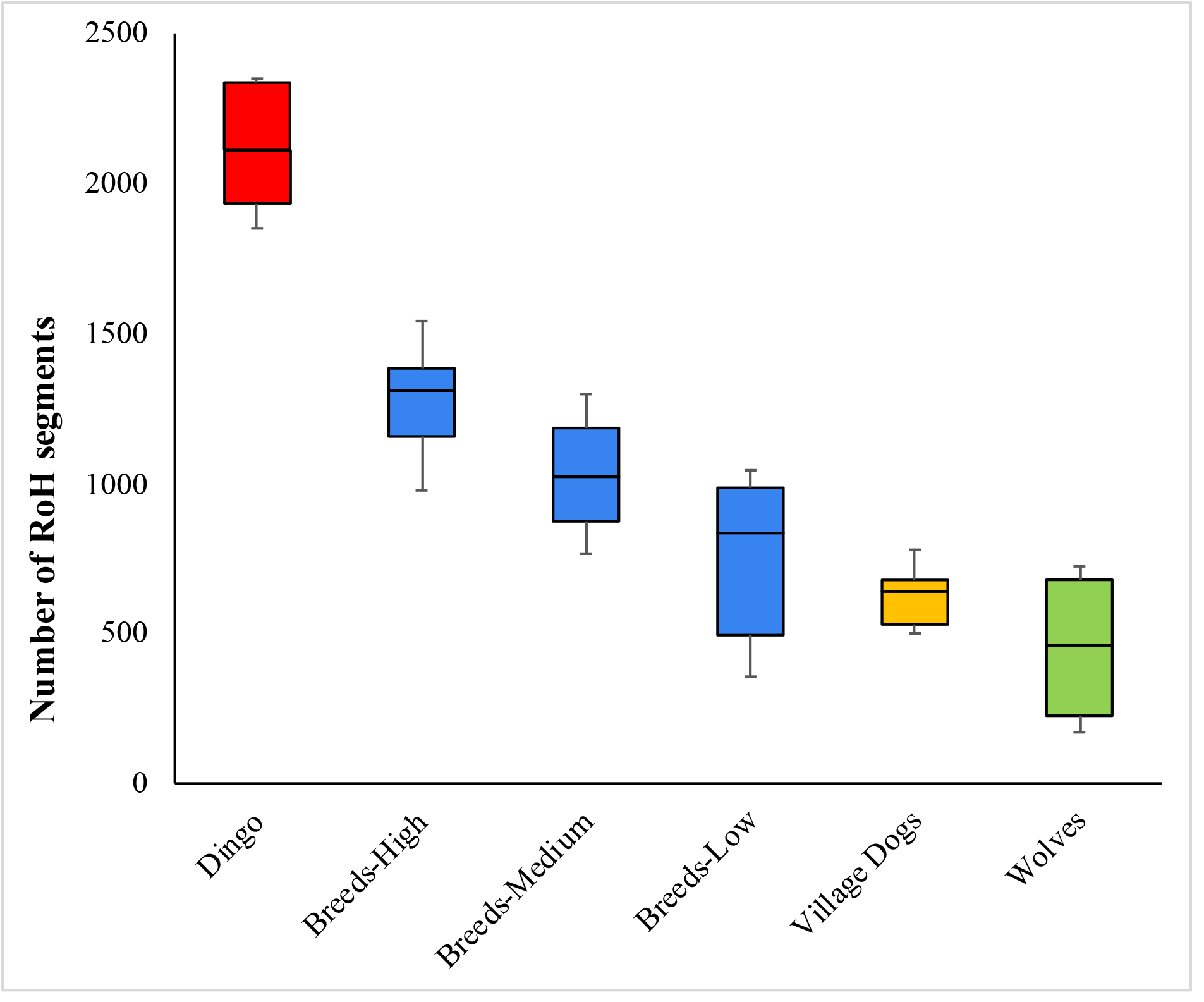
Box plot showing the mean estimates of the number of runs of homozygosity (RoH) segments for dingoes, dog breeds, village dogs, and wolves. A threshold of 0.5Mb or more was used to determine an RoH segment. The crosses indicate means, and the line within the box denotes the median. The whiskers show the maximum and minimum values.

### Deleterious mutational load in dingoes and other canines

To measure the accumulation of deleterious mutational load in canine genomes, we first estimated diversity at nonsynonymous (dN) and synonymous (dS) sites after identifying the protein-coding genes using genome annotations. The ratios (dN/dS) of the two estimates were then calculated, and those were plotted against the genomic diversities obtained for each genome. This analysis showed a highly significant negative correlation (*r* = 0.92, *P* < 0.000001) between the two variables (Figure 4). Importantly the dN/dS ratios of dingoes were predominantly higher than those of other canines, and the mean dN/dS estimate of dingoes (0.27) was significantly higher than that of highly inbred breed dogs (0.24) and other groups (*P* < 0.0001).

**Figure 4.**
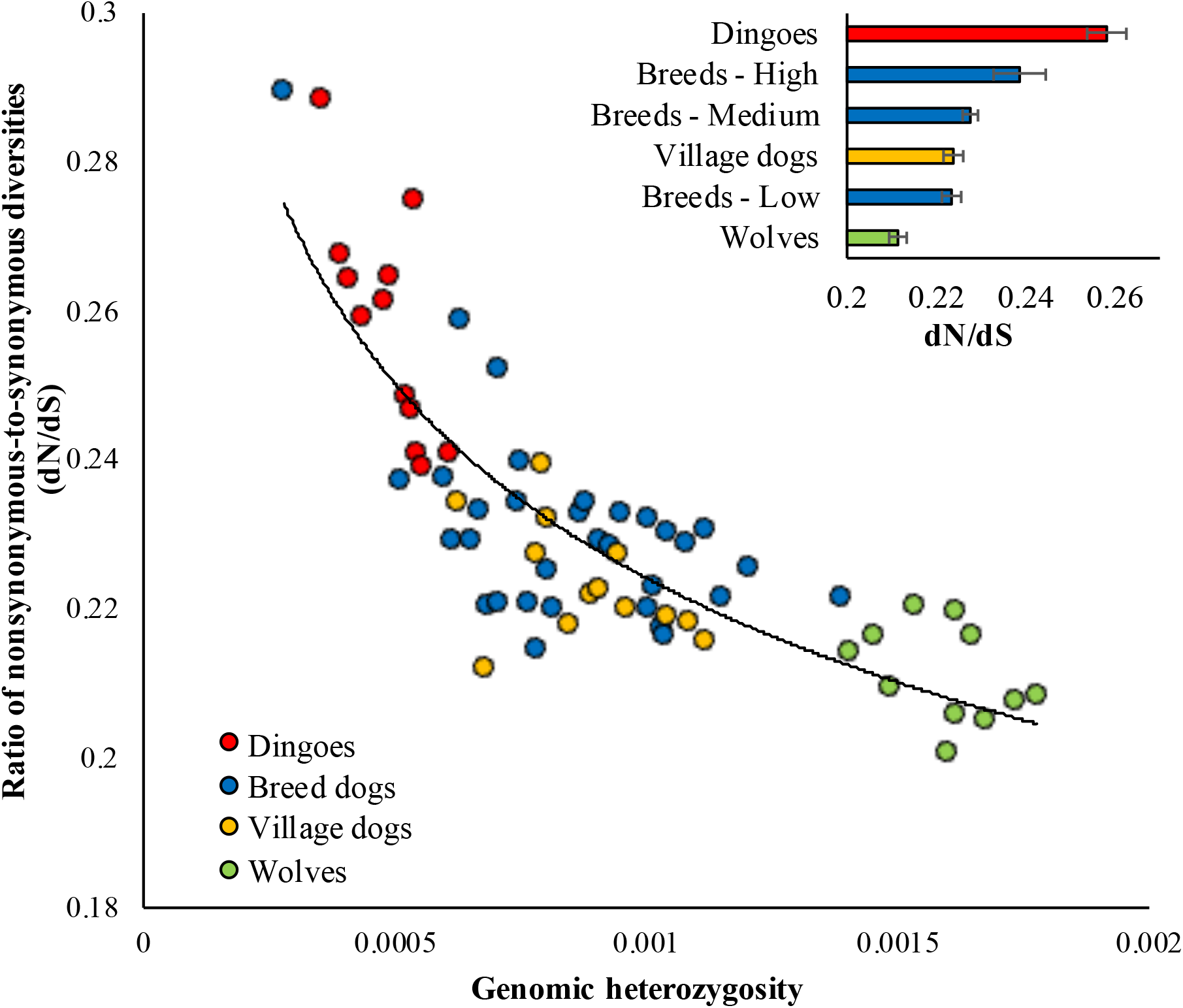
Correlation between the genomic diversity and the ratio of nonsynonymous-to-synonymous diversities (dN/dS). The relationship is highly significant (*r* = 0.92 and *P* < 0.00001). The best-fitting line is shown. **Inset**: Average dN/dS computed for various canine groups. Error bars show the standard error of the mean.

To measure the load of deleterious SNVs, we employed two methods. First, we used the SIFT scores to identify the deleterious nonsynonymous SNVs present in evolutionarily constrained positions. Second, the functional annotations were used to detect the highly deleterious LoF SNVs that cause premature termination of protein synthesis. Our analyses identified that dingoes had the highest load of homozygous deleterious missense SNVs and LoF SNVs of all the canines studied (Figure 5A). We observed that dingo genomes harboured an average of 1394 homozygous deleterious missense SNVs and 435 homozygous LoF SNVs, which were significantly (*P* < 0.0001) higher than those estimated for the highly inbred breed dogs (1239 and 406, respectively) and other canine groups. Wolves had the lowest number of homozygous deleterious missense SNVs (885) and homozygous LoF SNVs (283). We also plotted the proportions of homozygous and heterozygous deleterious (Figure 5B) and LoF SNVs (Figure 5C) in the canine genomes. This revealed that dingoes have the highest and wolves have the lowest proportion of homozygous SNVs. An opposite pattern was observed for the proportions of heterozygous SNVs in the canine genomes.

**Figure 5.**
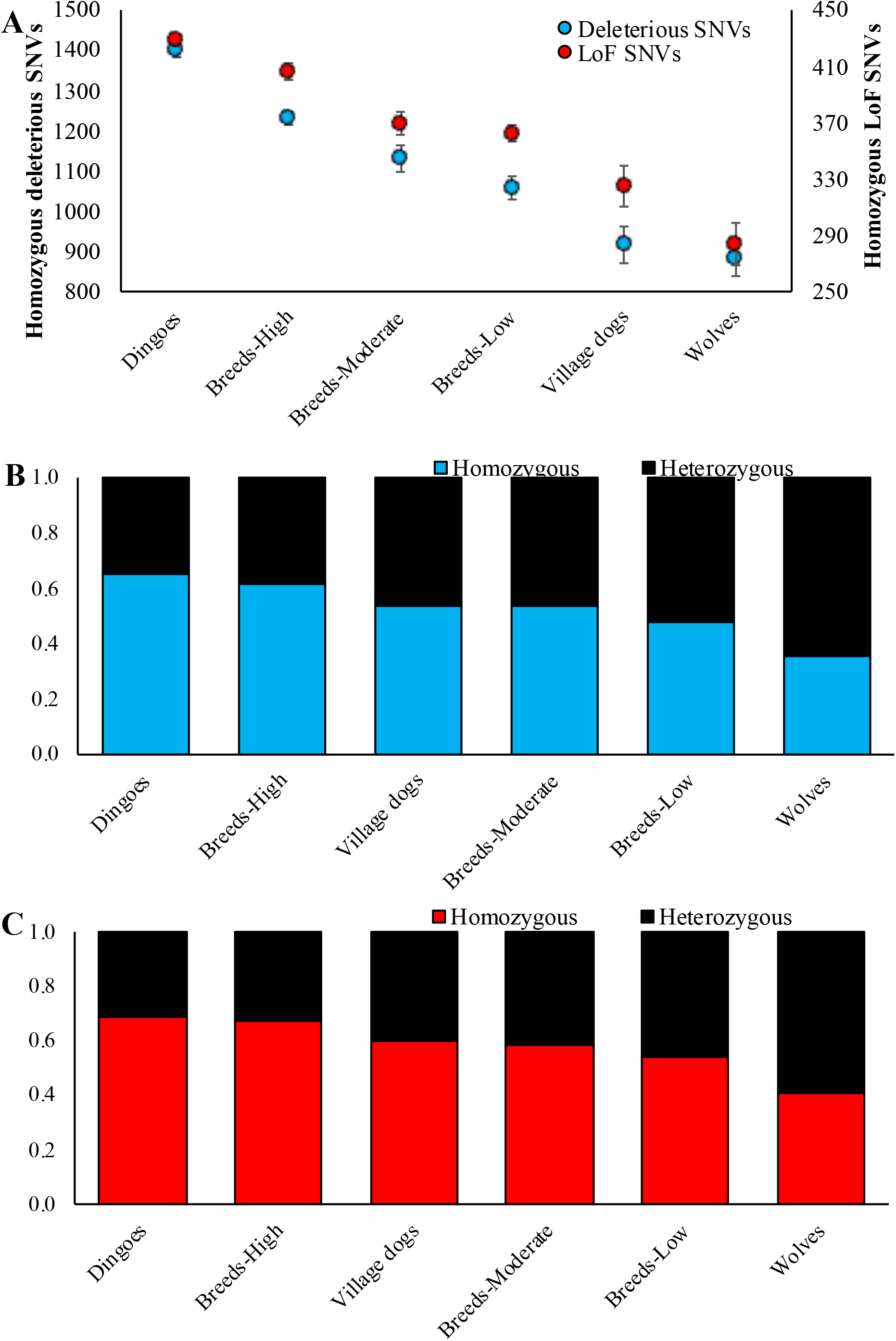
**(A)** Mean counts of deleterious and loss of function SNVs computed for the canine genomes. Error bars show the standard error of the mean. The mean estimates of homozygous deleterious SNVs and LoF SNVs of dingoes were statistically different from those of the highly inbred breeds and other groups (*P* < 0.0001). The stacked columns show the proportion of homozygous and heterozygous SNVs. **(B)** Deleterious SNVs **(C)** Loss of function SNVs.

## Discussion

Using whole-genome data, we examined the genomic signatures of dingoes, breed dogs, village dogs, and wolves. We found evidence for bottlenecks and founder effects in dingoes. Overall, despite the six-fold variation in genomic diversity observed within canines (0.0003-0.0018), dingoes had the lowest genomic diversity of all the canines studied (Figure 1). The variation in genomic diversity suggests that different canine populations may have large differences in effective population sizes. The mean diversity of dingoes was 36% less than that of highly inbred breed dogs and ∼4 times lower than that of wolves. The observed low diversity in dingoes corroborates previous reports using a small number of breeds (Freedman, et al. 2014; Zhang, et al. 2020). These findings have important implications for the conservation of dingoes but also for understanding their evolutionary history and relationship to other canines.

Previous studies showed that the dN/dS ratios for domesticated animals such as dog, cow, yak, pig, and silkworm and plants such as rice, soybean, and sunflower were significantly higher than those of their wild counterparts (Lu, et al. 2006; Mezmouk and Ross-Ibarra 2014; Renaut and Rieseberg 2015; Kono, et al. 2016; Marsden, et al. 2016; Pedersen, et al. 2017; Ramu, et al. 2017; Makino, et al. 2018; Peischl, et al. 2018; Xie, et al. 2018; Bosse, et al. 2019; Robinson, et al. 2019; Nicolas Dussex 2021; Subramanian 2021). Furthermore, domesticated plants and animals were found to have more homozygous deleterious SNVs than their wild relatives (Marsden, et al. 2016; Xie, et al. 2018; Subramanian 2021). In contrast, our results revealed the opposite: a much higher dN/dS ratio and more homozygous deleterious SNVs in wild dingoes than the domesticated breed dogs. Although dingoes have been in the wild for over thousands of years, their initial founder population must have been small. Theories predict that selection is not strong enough to remove deleterious mutations because the effect of genetic drift is high in small populations. A similar observation was reported for Przewalski horse population, which was almost extinct in 1969, and the ∼2,100 animals living now were derived from a very small captive stock of 12-15 animals (Orlando and Librado 2019). Genome analyses revealed a high dN/dS ratio and an elevated proportion of homozygous deleterious SNVs in this population.

Contemporary dingo populations are found across the entire continent, and hence the rate of historical inbreeding can be presumed to be low (due to the ongoing lethal control programs, contemporary populations in some regions are becoming small). Therefore, the observed low diversity and high load of deleterious mutations in dingoes could be due to a limited number of individuals in the initial founding populations. In contrast, the diversity and mutation load of breed dogs were modulated by intense inbreeding and artificial selection. To disentangle the bottleneck/founder effect from inbreeding, we examined the length of RoH segments in dingoes and dog breed genomes. Figure 3 shows the number of RoH that are >0.2 Mb. However, we also estimated the number of very long RoH that are >2 Mb. Overall, the dog breeds had 67 very long RoH segments, and dingoes had only 52 of such segments per genome, and this estimate for highly (100 per genome) and moderately inbred breeds (57 per genome) were higher than that of dingoes. It is well known that inbreeding leads to a few very long RoH (> 2Mb), and in contrast founder effect or bottleneck results in a greater number of medium-sized RoH (hundreds of Kbs) (Ceballos et a. 2018).

In this study, we found five lines of evidence for bottleneck and founder effects in dingo populations in Australia. First is the reduction in genomic heterozygosity in dingo populations, and second is the presence of a large number of RoH segments in the genomes of dingoes. We used three different methods to show the accumulation of harmful mutations in dingoes - the high dN/dS ratio, elevated number of homozygous deleterious amino acid changing SNVs, and LoF SNVs in dingo populations compared to breed dogs, village dogs, and wolves. These findings could potentially be explained by an initial bottleneck, which might have been caused by a limited number of founders that served as the ancestral stock for the modern dingoes. Similar results have been reported in other populations. For instance, the reduction in the heterozygosity of human populations negatively correlates with the geographical distance from Africa (DeGiorgio, et al. 2009). This is because when humans migrated out-of-Africa, only a small subpopulation served as the founders to the new location, and subsequent migrations from the new locations that are further away from Africa resulted in a serial founder effect (Henn, et al. 2015). These founder effects are also reflected in the deleterious mutation loads of the population, which positively correlate with geographical distance from Africa (Henn, et al. 2016). Similarly, populations that are far away from Africa also have a higher number of RoH than those in and close to Africa (Pemberton, et al. 2012). Genomes of the populations that migrated recently, such as French Canadians, also showed the signatures of founder effects, including low diversity, a large number of RoH segments, and elevated deleterious mutation load (Roy-Gagnon, et al. 2011). Genetic signatures of founder effects have also been reported in many other species, such as Isle Royale wolves (Robinson, et al. 2019), Corsican red deer (Hajji, et al. 2008), Afognak Island elk (Hundertmark and Van Daele 2010), and Common Myna (Hill and Pawley 2018), Wrangel Island mammoth(Rogers and Slatkin 2017), San Nicolas Island Island fox populations (Robinson, et al. 2016) and Steward Island kakapo populations (Dussex, et al. 2021).

The findings of this study help build our knowledge about the evolutionary history of dingoes and may have important implications for wildlife management. Our results highlight that despite their widespread distribution across Australia, the diversity of dingoes is still low compared to other canines, including the highly inbred dog breeds. We also found significant evidence of high accumulation of deleterious SNVs. Wildlife managers should begin considering the genomic consequences of lethal control programs on dingo populations. For example, aerial baiting programs which realise a 90% population knockdown (Ballard, et al. 2018) may initiate further bottlenecks and exacerbate the low genetic diversity and high mutational loads of targeted dingo populations with detrimental consequences. For example, low genomic diversity and mutational loads in dingoes may increase the threat of diseases such as Ehrlichiosis, Parvo virus, Sarcoptic mange, and Canine distemper to dingo populations. Further study of the genomic health of dingo populations at regional and local scales may be prudent to inform conservation and management practices. Ongoing genetic monitoring of the genetic diversity of lethally controlled populations may assist managers in designing conservation-aware dingo management plans, given the important ecological role and cultural significance of dingoes in the Australian landscape. The present investigation is based on a small number of dingoes, and hence there is a need for a wider study, including more genomes from various geographic locations. Such a study will also reveal any additional bottlenecks specific to different populations.

