## Supplementary data for "Genomic signatures of bottleneck and founder effects in dingoes"

**Supplementary Information**

**Figure S1: Locations of dingo samples**
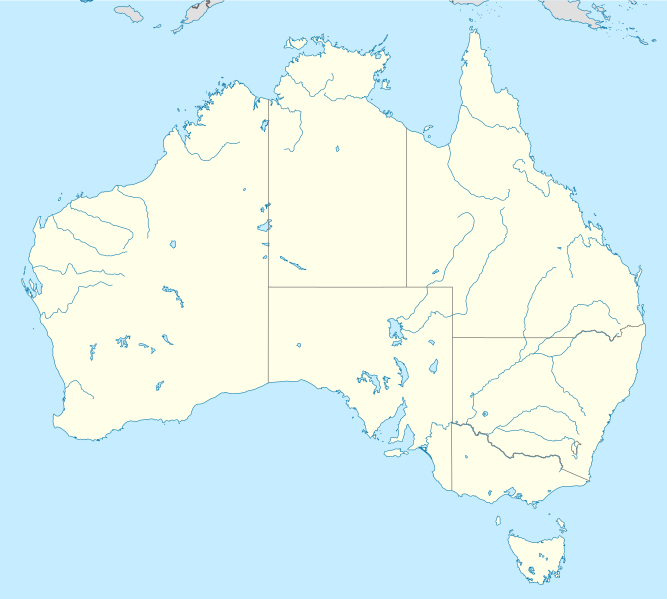

This study (one from

Zhang, et al (2020)

Admixed – not included

Note: One of the samples from Southeast Australia (blue circle in the bottom most corner) was obtained from captivity

**Table S1: Sequence Read Archive (SRA) accession numbers of the whole genomes of canines used in this study**

| **SRA ID** | **Canine**  **Population Name** | **Breed Name** | **Reference** |
| --- | --- | --- | --- |
| SRR7107643 | Breeds-High | Norwegian Lundehund | [1] |
| SRR7120152 | Breeds-High | Bull Terrier | [1] |
| SRR7107885 | Breeds-High | Basenji | [1] |
| SRR7107902 | Breeds-High | Soft Coated Wheaten Terrier | [1] |
| SRR7107968 | Breeds-High | Border Terrier | [1] |
| SRR7120213 | Breeds-High | Scottish Terrier | [1] |
| SRR7107598 | Breeds-High | Entlebucher Sennenhund | [1] |
| SRR7107922 | Breeds-High | Airedale Terrier | [1] |
| SRR7107634 | Breeds-High | Weimaraner | [1] |
| SRR7107973 | Breeds-High | Berger Picard | [1] |
| SRR7120170 | Breeds-High | Irish Water Spaniel | [1] |
| SRR7107963 | Breeds-High | Field Spaniel | [1] |
| SRR7107867 | Breeds-High | Australian Cattle Dog | [1] |
| SRR7107883 | Breeds-High | English Springer Spaniel | [1] |
| SRR7107578 | Breeds-Low | Border Collie | [1] |
| SRR7107891 | Breeds-Low | Labrador Retriever | [1] |
| SRR2094392 | Breeds-Low | Chow Chow | [1] |
| SRR7107916 | Breeds-Low | Yorkshire Terrier | [1] |
| SRR7107838 | Breeds-Low | Peruvian Inca Orchid | [1] |
| SRR7107992 | Breeds-Low | Alaskan Malamute | [1] |
| SRR7107895 | Breeds-Low | Tibetan Terrier | [1] |
| SRR2095503 | Breeds-Low | Saluki | [1] |
| SRR2095478 | Breeds-Low | Chihuahua | [1] |
| SRR2095539 | Breeds-Low | Siberian Husky | [1] |
| SRR7120156 | Breeds-Low | Carolina Dog | [1] |
| SRR7107795 | Breeds-Moderate | Greyhound | [1] |
| SRR7107657 | Breeds-Moderate | Afghan Hound | [1] |
| SRR7107839 | Breeds-Moderate | Swedish Lapphund | [1] |
| SRR7107884 | Breeds-Moderate | German Shepherd Dog | [1] |
| SRR7107933 | Breeds-Moderate | Shiba Inu | [1] |
| SRR5664959 | Breeds-Moderate | Cocker Spaniel (American) | [1] |
| SRR7120187 | Breeds-Moderate | Miniature Poodle | [1] |
| SRX7276152 | Dingo | Fraser Dingo | [2] |
| Y3 | Dingo | Fraser Dingo | This study |
| Y4 | Dingo | Fraser Dingo | This study |
| B44 | Dingo | Fraser Dingo | This study |
| Y6 | Dingo | Fraser Dingo | This study |
| Y2 | Dingo | Fraser Dingo | This study |
| SRX7276159 | Dingo | Dingo-NW | [2] |
| SRX7276160 | Dingo | Dingo-NW | [2] |
| SRX7276154 | Dingo | Dingo-NW | [2] |
| SRX7276158 | Dingo | Dingo-NW | [2] |
| SRX7276153 | Dingo | Dingo-NW | [2] |
| SRX7276156 | Dingo | Dingo-NW | [2] |
| Ernie | Dingo | Dingo-NW | This study |
| Kimmi | Dingo | Dingo-NW | This study |
| Mikey | Dingo | Dingo-SE | This study |
| SRX7276151 | Dingo | Dingo-SE | [2] |
| SRX7276157 | Dingo | Dingo-SE | [2] |
| SRX7276155 | Dingo | Dingo-SE | [2] |
| SRR7107676 | Village Dog | Village Dog - Egypt | [1] |
| SRR7107649 | Village Dog | Village Dog - China | [1] |
| SRR2095463 | Village Dog | Village Dog – Bernese Mountain Dog | [1] |
| SRR7107690 | Village Dog | Village Dog - Lebanon | [1] |
| SRR7107703 | Village Dog | Village Dog - Vietnam | [1] |
| SRR7107828 | Village Dog | Nigerian Indigenous Dog | [1] |
| SRR7107684 | Village Dog | Village Dog - India | [1] |
| SRR7107700 | Village Dog | Village Dog - Qatar | [1] |
| SRR7107697 | Village Dog | Village Dog - Papua New Guinea | [1] |
| SRR7107693 | Village Dog | Village Dog - Namibia | [1] |
| SRR7107698 | Village Dog | Village Dog - Portugal | [1] |
| SRR7107823 | Village Dog | Vietnamese Indigenous Dog | [1] |
| SRR7107702 | Village Dog | Village Dog - Taiwan | [1] |
| SRR7107786 | Wolf | Wolf | [1] |
| SRR7107910 | Wolf | Wolf | [1] |
| SRR7107787 | Wolf | Wolf | [1] |
| SRR7107542 | Wolf | Wolf | [1] |
| SRR7107909 | Wolf | Wolf | [1] |
| SRR7107540 | Wolf | Wolf | [1] |
| SRR7107783 | Wolf | Wolf | [1] |
| SRR7107776 | Wolf | Wolf | [1] |
| SRR7107778 | Wolf | Wolf | [1] |
| SRR7107987 | Wolf | Wolf | [1] |
| SRR7107777 | Wolf | Wolf | [1] |
